# Cardiac atrophy, dysfunction, and metabolic impairments: a cancer-induced heart failure phenotype

**DOI:** 10.1101/2023.09.30.560250

**Authors:** Leslie M. Ogilvie, Luca J. Delfinis, Bridget Coyle-Asbil, Vignesh Vudatha, Razan Alshamali, Bianca Garlisi, Madison Pereira, Kathy Matuszewska, Madison C. Garibotti, Shivam Gandhi, Keith R. Brunt, Jose G. Trevino, Christopher G.R. Perry, Jim Petrik, Jeremy A. Simpson

## Abstract

Muscle atrophy and weakness are prevalent features of cancer. While extensive research has characterized skeletal muscle wasting in cancer cachexia, limited studies have investigated how cardiac structure and function are affected by therapy-naïve cancer. In cell-based models of orthotopic, syngeneic epithelial ovarian cancer (EOC) and pancreatic ductal adenocarcinoma (PDAC), and a patient-derived pancreatic xenograft model (PDX), we evaluated cardiac structure, function, and metabolism. Tumor-bearing mice showed cardiac atrophy and intrinsic systolic and diastolic dysfunction; associated with hypotension and exercise intolerance. In hearts of ovarian tumor-bearing mice, fatty acid-supported mitochondrial respiration decreased and carbohydrate-supported respiration increased, establishing a substrate shift in cardiac metabolism that is characteristic of heart failure. EOC decreased cytoskeletal and cardioprotective gene expression, which was paralleled by downregulation of transcription factors that regulate cardiomyocyte size and function. PDX tumors altered myosin heavy chain isoform expression – a molecular phenotype observed in heart failure. Markers of autophagy and ubiquitin-proteasome system were upregulated with cancer, providing evidence of catabolic signaling that promotes cardiac wasting. Together, metabolic stress, cardiac gene dysregulation, and upregulation of catabolic pathways contribute to cardiac atrophy and failure during cancer. Finally, we demonstrate that pathological cardiac remodeling is induced by human cancer, providing translational evidence of cancer-induced cardiomyopathy.

## INTRODUCTION

Cancer and cardiovascular disease (CVD) are the two leading causes of death worldwide.^1,2^ Although generally thought of as distinct diseases, intersections between cancer and CVD have emerged in the past decade. The field of cardio-oncology has developed with the aim of improving the identification, monitoring, and treatment of cardiovascular complications in cancer patients during and after cancer therapy. Research has largely focused on how chemotherapeutics impair cardiovascular function.^3^ Indeed, the cumulative effects of cardiotoxic therapy and the presence of CVD risk factors leads to long-term morbidity and poor quality of life in cancer patients, even when cured of cancer.^4^ Recent evidence suggests that cancer itself, in the absence of exposure to cardiotoxic therapeutics, impacts negatively upon cardiovascular health.^5^ However, our understanding of the cardiac sequelae of cancer, in a therapy-naïve setting, requires further investigation.

Cancer cachexia, affecting approximately 50-80% of cancer patients, is a complex metabolic syndrome, characterized by the progressive loss of body mass, predominantly from skeletal muscle and fat, and multiple organs including the liver, kidney, spleen, gastrointestinal tract, and heart.^6,7^ Skeletal muscle weakness and atrophy are well-known consequences of cancer cachexia.^8^ Muscle wasting is attributed to a decline in protein synthesis and an increase in degradation, mediated by systemic increases in inflammatory cytokines released by activated immune cells of the tumor and host.^9^ These cytokines promote hyperactivation of ubiquitin-proteasome system (UPS)-mediated protein degradation, leading to the breakdown of myofibrillar proteins and muscle atrophy.^7^ Recent evidence demonstrated that autophagic flux is upregulated in conditions of nutrient or growth factor deprivation and contributes to skeletal muscle depletion in cancer cachexia.^10,11^ Subsequently, proteins and organelles are targeted for lysosomal degradation and the resulting molecular components are recycled as substrates to support metabolic demand. Although both autophagic and UPS activation are implicated in the development of skeletal muscle wasting in cancer, the contribution of these pathways to cancer-induced cardiac atrophy is debated.

At the molecular level, cardiac transcription factors regulate the expression of genes encoding structural and regulatory proteins in the myocardium.^12^ Several transcription factors – GATA binding protein (GATA), Myocyte enhancing factor 2 (MEF2), Nuclear factor of activated T cells (NFAT) and Serum response factor (SRF) – regulate the cardiac gene program in embryonic development and in response to mitogenic or hemodynamic stimuli.^13^ The upregulation of these transcription factors in adults elicits hypertrophic growth, however, each factor is differentially regulated by varying stimuli. Thus, how cancer alters the expression of cardiac transcription factors in atrophied hearts remains unknown.

We assessed cardiac morphology and function in murine implanted and patient xenografted cancer models: 1) cell-based epithelial ovarian cancer (EOC), 2) cell-based pancreatic ductal adenocarcinoma (PDAC), and 3) patient-derived pancreatic cancer xenograft (PDX). Clinically, cachexia is a frequent and prominent feature of these cancer types.^14,15^ We also determined whether cancer perturbed cardiac substrate metabolism and increased oxidative stress levels in the myocardium, given such stress responses have been reported in skeletal muscle during cancer.^8,16,17^ To investigate the underlying mechanisms of cancer-induced cardiac impairments, we quantified the cardiac gene program and catabolic pathways that regulate protein degradation in the heart. We show that cancer causes cardiac atrophy, dysfunction, metabolic dysregulation, and exercise intolerance. Further, we provide novel insight into the key pathways that drive cardiac wasting and eventual failure.

## RESULTS

### Mice with ovarian cancer show impaired exercise tolerance and reduced skeletal muscle mass

Tumors were induced as orthotopic models of ovarian or pancreatic cancer. We first assessed the characteristics of advanced ovarian cancer by measuring morphometrics and voluntary exercise capacity. Mice with EOC presented with large primary tumors, extensive secondary peritoneal lesions, a loss of body mass, and anemia (Figure 1A-C; Table S2). This was associated with a profound decrease in the mass of tibialis anterior, extensor digitorum longus, and soleus muscles (Figure 1D; Table S2) and exercise intolerance (voluntary running distance; Figure 1E). These data confirm a cachectic phenotype that is prevalent in advanced ovarian and pancreatic cancers.

**Figure 1.**
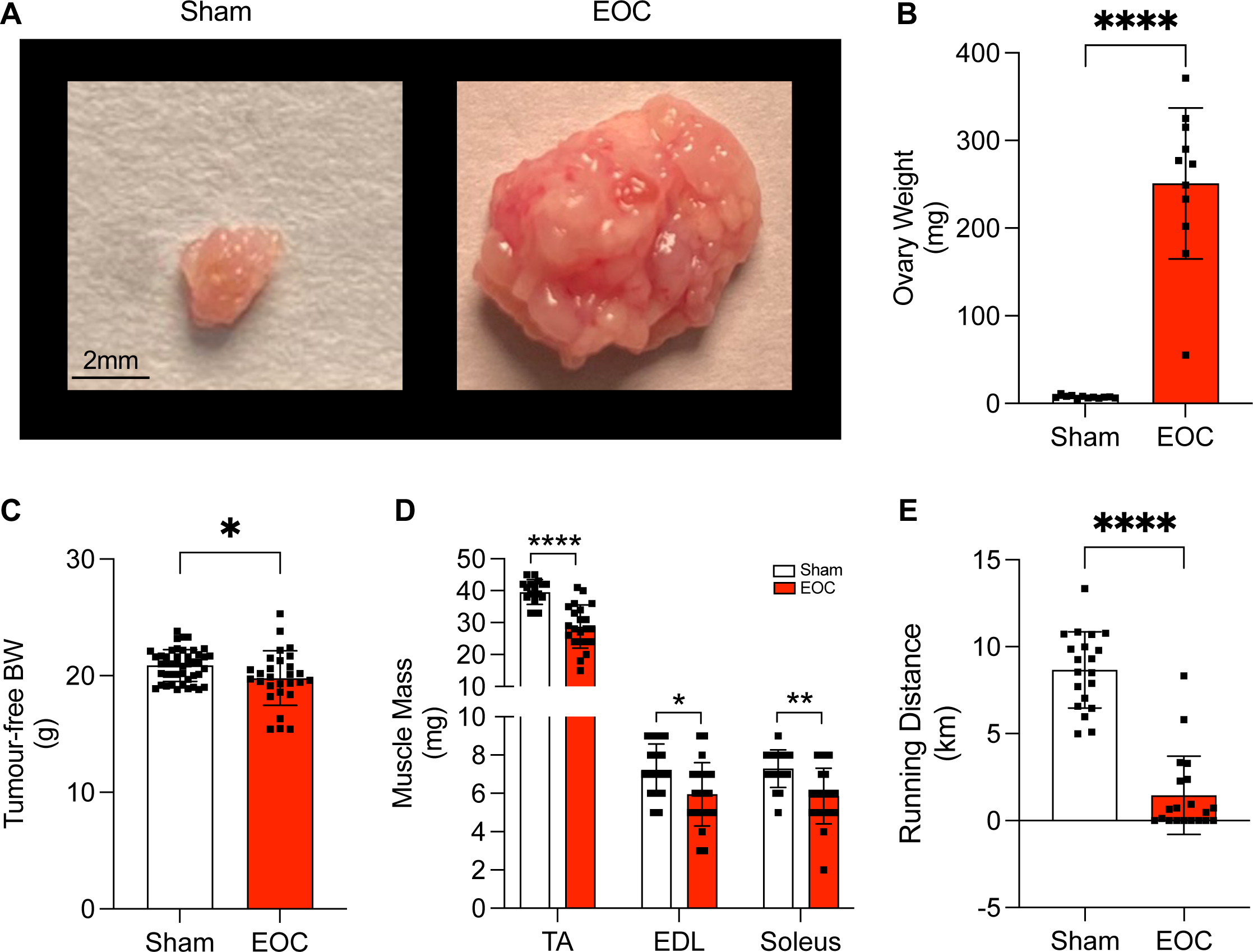
Phenotypic characterization of a mouse model of advanced ovarian cancer. (A) Representative images of ovaries from sham mice and tumors from EOC mice. (B) Ovary weight. (C) Tumor-free body weight (BW). (D) Skeletal muscle mass (TA, tibialis anterior; EDL, extensor digitorum longus). (E) Average voluntary running distance. Data are means ± SD. Significance determined by an unpaired student’s t-test.

### Ovarian cancer causes cardiac atrophy

Ovarian tumor-bearing mice showed a ∼10% decrease in heart mass and reduced left ventricle, right ventricle, and atrial mass (Figure 2A-C; Table 1), confirming that this is not a ventricle-specific response, but a global atrophy of all chambers. To corroborate that cancer alters cardiac structure, we performed echocardiography on sham and EOC mice (Figure 2D; Table 1). Mice with ovarian cancer showed no changes in end systolic dimension (ESD), end diastolic dimension (EDD; Figure 2E), ejection fraction (Figure 2F), or fraction shortening (Table 1). Cardiac output decreased in tumor-bearing mice, likely due to non-significant decreases in heart rate and stroke volume (Figure 2G; Table 1). Consistent with the observed decrease in heart mass, posterior wall thickness decreased in tumor-bearing mice (Figure 2H), providing further evidence of cardiac atrophy.

**Figure 2.**
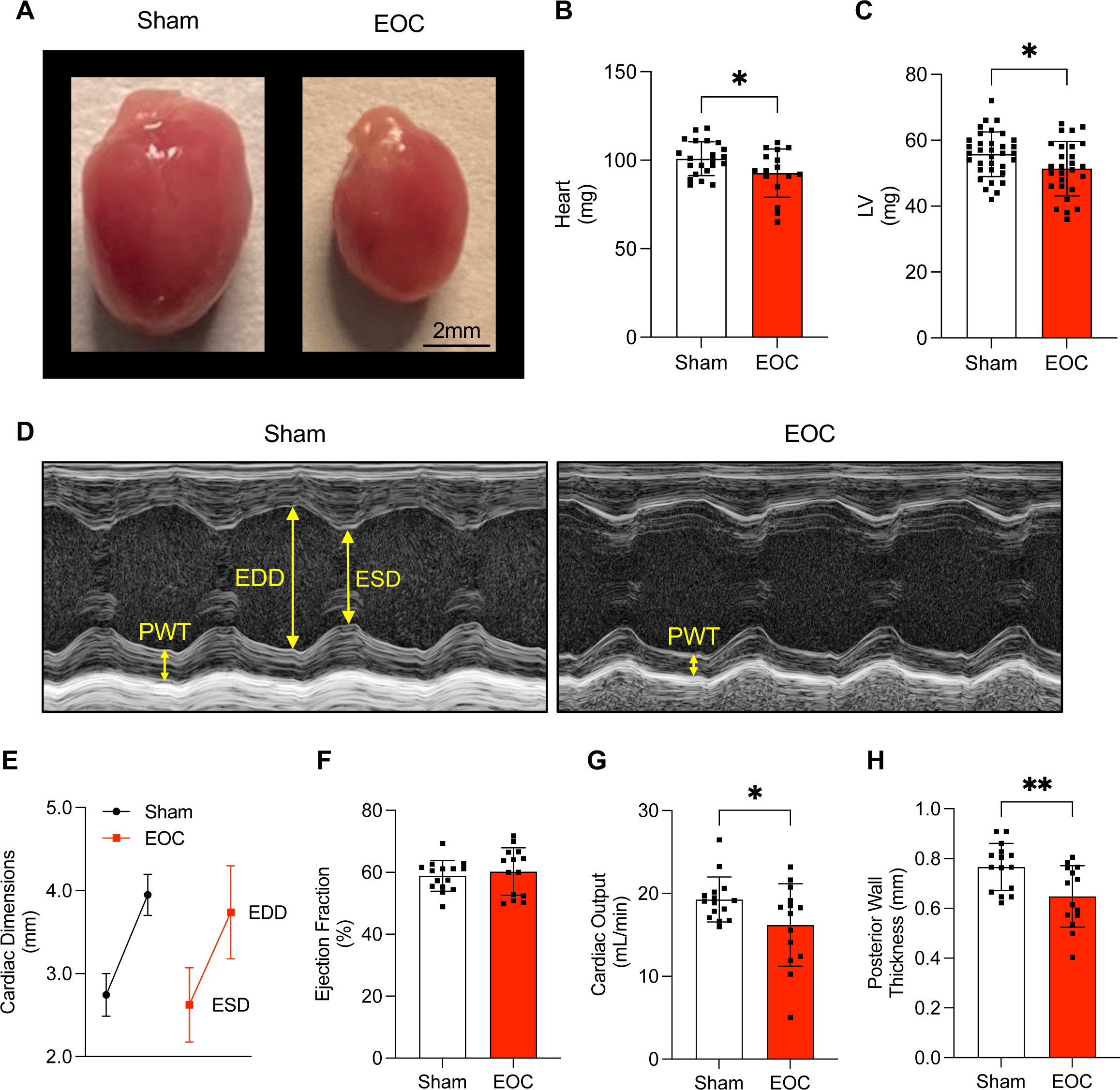
Comparison of cardiac parameters by echocardiography. (A) Representative gross morphology of hearts from sham and EOC mice. (B) Heart mass. (C) LV mass. (D) Representative echocardiogram of the LV from sham and EOC mice. (E) Cardiac dimensions; EDD, end diastolic dimension (top); ESD, end systolic dimension (bottom). (F) Ejection fraction. (G) Cardiac output. (H) LV posterior wall thickness. Data are means ± SD. Significance determined by an unpaired student’s t-test.

**Table 1.** Comparison of cardiac structural and functional parameters between sham and ovarian cancer mice.

|  | Sham | EOC | p-value (summary) |
| --- | --- | --- | --- |
| *Morphometric Parameters*<br>(n=16-36) |  |  |  |
| Heart (mg) | 100.9 ± 9.6 | 92.3 ± 13.6 | 0.036 (*) |
| LV (mg) | 55.7 ± 6.8 | 51.4 ± 8.3 | 0.027 (*) |
| RV (mg) | 17.7 ± 2.0 | 16.2 ± 3.5 | 0.040 (*) |
| Atria (mg) | 5.4 ± 0.9 | 4.7 ± 1.0 | 0.009 (**) |
| *Echocardiographic Parameters*<br>(n=14-15) |  |  |  |
| End systolic dimension (mm) | 2.7 ± 0.3 | 2.6 ± 0.4 | 0.381 (ns) |
| End diastolic dimension (mm) | 4.0 ± 0.2 | 3.8 ± 0.5 | 0.196 (ns) |
| Posterior wall thickness (mm) | 0.8 ± 0.10 | 0.6 ± 0.12 | 0.007 (**) |
| Heart rate (bpm) | 477 ± 27 | 465 ± 53 | 0.475 (ns) |
| Stroke volume (μL/beat) | 40.5 ± 5.4 | 37.4 ± 10.1 | 0.539 (ns) |
| Cardiac output (mL/min) | 19.3 ± 2.7 | 16.2 ± 5.0 | 0.046 (*) |
| Ejection fraction (%) | 58.9 ± 4.9 | 60.2 ± 7.7 | 0.566 (ns) |
| Fractional shortening (%) | 30.2 ± 3.9 | 31.2 ± 4.7 | 0.524 (ns) |
| *Invasive Hemodynamic Parameters*<br>(n=14-17) |  |  |  |
| LVP Max (mmHg) | 98.0 ± 2.3 | 78.8 ± 11.1 | <0.0001 (****) |
| End diastolic pressure (mmHg) | 3.5 ± 1.7 | 2.7 ± 2.1 | 0.198 (ns) |
| dP/dt Max (mmHg/s) | 9462 ± 870 | 7122 ± 1996 | 0.0004 (***) |
| dP/dt @LVP40 (mmHg/s) | 8669 ± 893 | 6825 ± 2029 | 0.003 (**) |
| dP/dt Min (mmHg/s) | -9647 ± 903 | -7250 ± 1772 | <0.0001 (****) |
| Tau (Glantz) (ms) | 8.2 ± 0.7 | 7.7 ± 1.3 | 0.171 (ns) |
| Heart rate (bpm) | 502 ± 40 | 456 ± 62 | 0.023 (*) |
| Systolic BP (mmHg) | 95.6 ± 4.3 | 73.0 ± 11.9 | <0.0001 (****) |
| Diastolic BP (mmHg) | 67.0 ± 4.8 | 40.0 ± 14.0 | <0.0001 (****) |
| MAP (mmHg) | 76.5 ± 4.6 | 50.2 ± 12.4 | <0.0001 (****) |
| *Langendorff Parameters*<br>(n=3-4) |  |  |  |
| Heart rate (bpm) | 448 ± 0.6 | 446 ± 0.0 | 0.0019 (**) |
| LVP Max (mmHg) | 101.3 ± 13.8 | 53.3 ± 14.0 | 0.0062 (**) |
| End diastolic pressure (mmHg) | 8.0 ± 1.0 | 8.3 ± 0.5 | >0.999 (ns) |
| dP/dt Max (mmHg/s) | 4629 ± 890 | 2087 ± 537 | 0.0051 (**) |
| dP/dt Min (mmHg/s) | -2802 ± 329 | -1257 ± 383 | 0.0025 (**) |
| Tau (Weiss) (ms) | 12.7 ± 1.7 | 17.0 ± 2.9 | 0.0734 (ns) |

Values are mean ± SD. EOC, epithelial ovarian cancer; LV, left ventricle; RV, right ventricle; bpm, beats per minute; LVP Max, maximum left ventricle pressure; dP/dt Max, maximum rate of change of pressure during systole; dP/dt Min, maximum negative rate of change of pressure during diastole; Tau, relaxation time constant; BP, blood pressure; MAP, mean arterial pressure. Significance determined by an unpaired student's t-test; ns, not significant. Mann-Whitney test used on non-normally distributed data, where appropriate.

### Ovarian cancer impairs cardiac function, independent of external factors

Next, we performed invasive hemodynamics to provide a highly sensitive assessment of systolic and diastolic function of the heart (Figure 3A; Table 1). Tumor-bearing mice showed a decrease in heart rate (Figure 3B) and maximum LV pressure (Figure 3C). Maximal rates of contraction (dP/dt Max) and relaxation (dP/dt Min) decreased compared to shams (Figure 3D), with no change in passive filling pressure (EDP) or the relaxation time constant, Tau (Table 1). Tumor-bearing mice presented with hypotension, decreases in mean arterial pressure (MAP; Figure 3E), systolic (SBP), and diastolic blood pressure (DBP; Figure 3F). These data provide new evidence that ovarian cancer has detrimental effects on cardiac contraction, relaxation, and blood pressure regulation directly.

**Figure 3.**
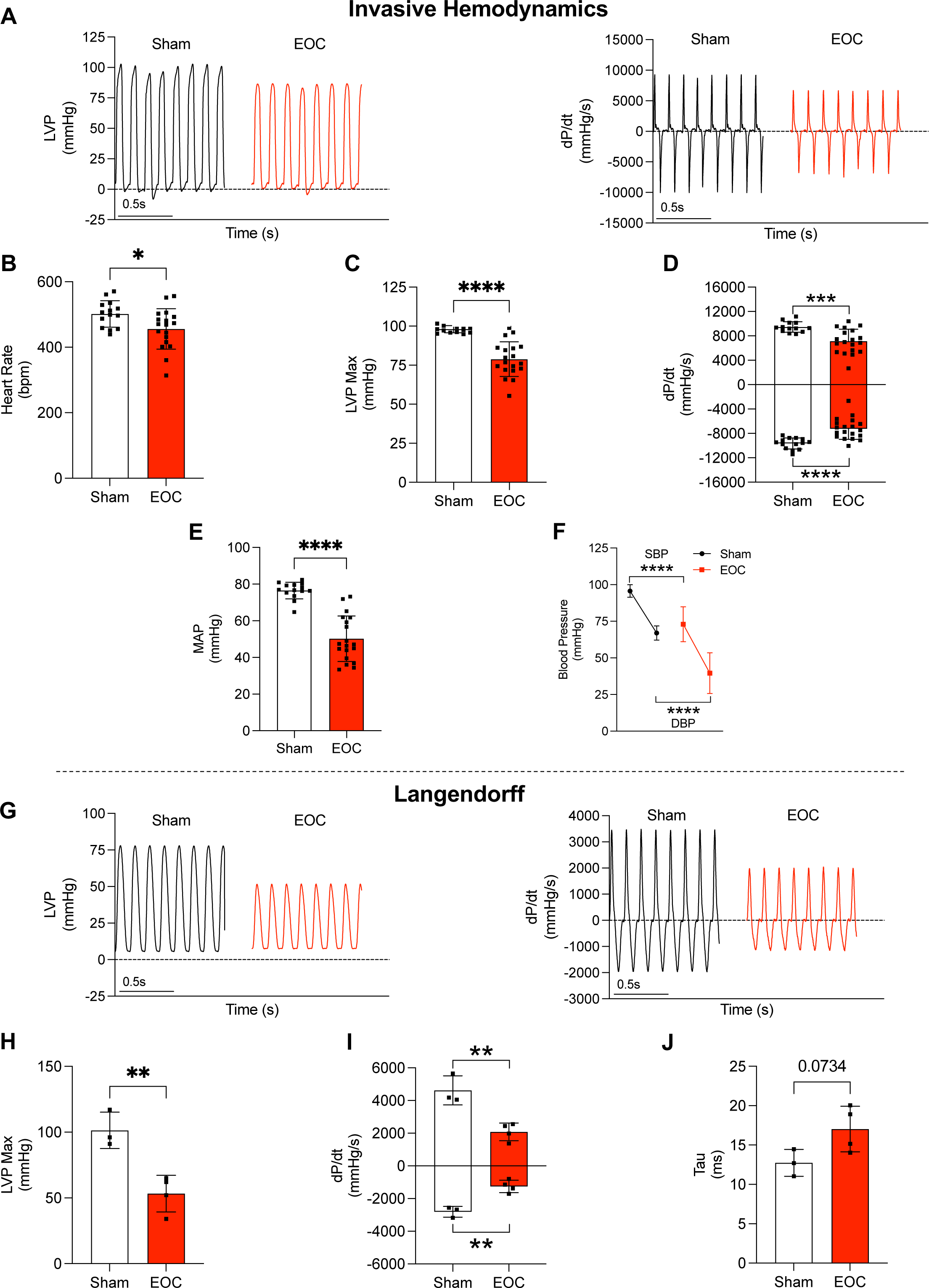
Evaluation of cardiac function by invasive hemodynamics and Langendorff. (A) Representative tracings of left ventricular pressure (LVP; left) and rate of change of pressure (dP/dt; right) by invasive hemodynamics from sham and EOC mice. (B) Heart rate. (C) Maximum LVP. (D) Maximum (dP/dt Max; top) and minimum (dP/dt Min; bottom) rate of change of pressure. (E) Mean arterial pressure (MAP). (F) Systolic (SBP; top) and diastolic blood pressure (DBP; bottom). (G) Representative tracings of LVP (left) and dP/dt (right) by Langendorff from sham and EOC mice. (H) Maximum LVP. (I) dP/dt Max (top) and Min (bottom). (J) Relaxation time constant (Tau). Data are means ± SD. Significance determined by an unpaired student’s t-test.

The *in vivo* assessment of cardiac function is dependent on preload, afterload, and heart rate.^18^ In situations where these parameters are not equivalent (e.g., peripheral fluid shifting due to cancer), measures of intrinsic cardiac function can be misinterpreted. Thus, to validate that cardiac dysfunction is independent of external influences (e.g., changes in hemodynamic load, neurohormonal input), we performed Langendorff *ex vivo* isolated heart assessments (Figure 3G, Table 1). Maximal LV pressure decreased in EOC mice (Figure 3H). Peak contraction (dP/dt Max) and relaxation rates (dP/dt Min) decreased by ∼55% compared to shams (Figure 3I), and the relaxation time constant (Tau) tended to increase (p=0.07; Figure 3J). Therefore, we concluded that cancer-induced systolic and diastolic dysfunction are intrinsic to the myocardium, which persists independent of central neurohormonal influences or changes in hemodynamic loading.

### Cardiac metabolism substrate shift and altered mH_2_O_2_ due ovarian cancer

Mitochondrial dysfunction is implicated in the development of skeletal muscle dysfunction and atrophy in cancer cachexia.^8^ Thus, we evaluated if mitochondrial oxygen consumption was impaired in the LV of ovarian tumor-bearing mice as an index of ATP synthesis by oxidative phosphorylation. We used pyruvate and malate to mimic mitochondrial respiration under carbohydrate-supported conditions and L-carnitine with palmitoyl-CoA to mimic fat-supported conditions. This assay design allowed us to determine if changes in oxidative phosphorylation differed between macronutrients. ADP-stimulated, pyruvate/malate-supported mitochondrial respiration increased in EOC mice at every ADP concentration (Figure 4A; main effects of cancer and ADP concentration, interaction effect). Conversely, ADP-stimulated, L-carnitine/palmitoyl-CoA-supported mitochondrial respiration showed that oxygen consumption decreased in the LV of ovarian tumor-bearing mice at all ADP concentrations (Figure 4B; main effect of cancer). These changes in mitochondrial oxygen consumption were not explained by differences in ETC subunit contents as there was no change in tumor-bearing mice relative to shams (Figure 4C, S1A). These metabolic alterations are consistent with a heart failure phenotype, where substrate oxidation for ATP production is shifted from heavy reliance on fatty acids to greater carbohydrate oxidation.^19^ This concept was further supported when we evaluated pyruvate/malate ADP-stimulated state II respiration (Figure S1B), suggesting even pyruvate supported uncoupled respiration is elevated as well as the additional stimulatory effect of coupled glutamate oxidation (state III Δ glutamate respiration; Figure S1C), and state III Δ succinate respiration (Figure S1D) as the LV of EOC mice exhibited an increase under all conditions. Conversely, no significant changes were seen under L-carnitine/Palmitoyl CoA state II respiration (Figure S1E) or state III Δ succinate (Figure S1F).

**Figure 4.**
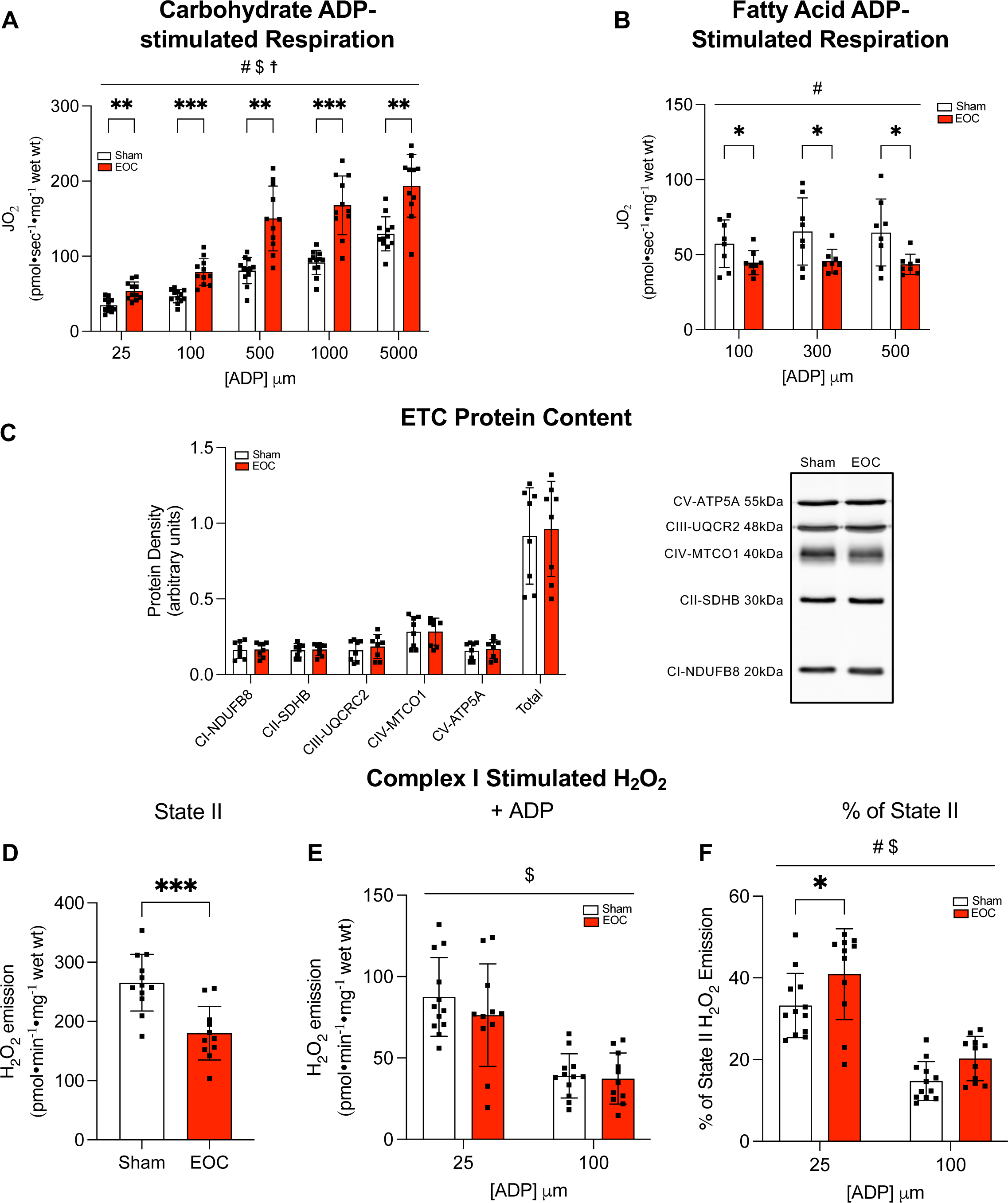
Altered mitochondrial oxygen consumption and mH_2_O_2_ emission in tumor-bearing mice. (A) Carbohydrate ADP-stimulated respiration. (B) Fat ADP-stimulated respiration. (C) Protein content of electron transport chain (ETC) subunits (left) and representative western blot for ETC content in sham and EOC mice (right). (D) mH_2_O_2_ emission supported by pyruvate (10mM) and malate (2 mM) (NADH) assessed under maximal state II (no ADP) conditions. (E) mH_2_O_2_ emission across a range of [ADP] to model metabolic demand. (F) mH_2_O_2_ emission expressed as a % of maximal state II levels. Data are means ± SD. A, B, E, F determined by two-way ANOVA. # main effect of cancer, $ main effect of ADP concentration, IZ interaction effect, * indicates significant difference between sham and EOC. C, F determined by unpaired student’s t-tests.

We next stimulated complex I with pyruvate and malate to generate NADH in the absence of ADP to elicit maximal mH_2_O_2_ emission, a type of mitochondrial reactive oxygen species (ROS). We then introduced varying concentrations of ADP into the assay medium to determine the ability of ADP to attenuate this emission. In doing so, the degree of mH_2_O_2_ could be assessed in the physiological relevant context of ATP synthesis.^20^ Maximal mH_2_O_2_ emission rates were decreased in the LV of EOC mice (Figure 4D), however, there was no difference in the ability of cardiomyocytes to attenuate mH_2_O_2_ with the introduction of ADP (Figure 4E; main effect of ADP concentration). Interestingly, when ADP-suppressed mH_2_O_2_ emission was expressed as a percentage of total mH_2_O_2_ emission, tumor-bearing mice showed a relative increase in mH_2_O_2_ (Figure 4F; main effects of cancer and ADP concentration). This suggests EOC mice generate less mH_2_O_2_ under maximal conditions (absence of ADP or ATP synthesis), yet the ability of ADP to attenuate mH_2_O_2_ suggests a reduced mitochondrial responsiveness to ADP’s suppressive effect on mH_2_O_2_. While such recalibration of ADP’s control of mH_2_O_2_ is perplexing, the similar absolute rate during actual ATP synthesis in EOC mice suggests a dynamic remodeling that might serve to maintain net mH_2_O_2_ during this stage of cancer. Overall, these data provide new insight that ovarian cancer causes metabolic stress in the heart, demonstrated by a substrate shift towards heightened carbohydrate-supported metabolism.

### Tumor-bearing mice show pathological remodeling of the left ventricle

To evaluate cardiac morphology on a cellular level, we quantified cardiomyocyte cross-sectional area (CSA), capillary density, and fibrosis by histology (Figure 5A). CSA decreased by ∼25% in ovarian tumor-bearing mice, as compared to shams, indicating an atrophic phenotype of the heart (Figure 5B, C). Cardiac atrophy was associated with a trending decrease in cardiomyocyte capillary density (p=0.07; Figure 5D), but no change in the quantity of interstitial fibrosis (Figure S2A).

**Figure 5.**
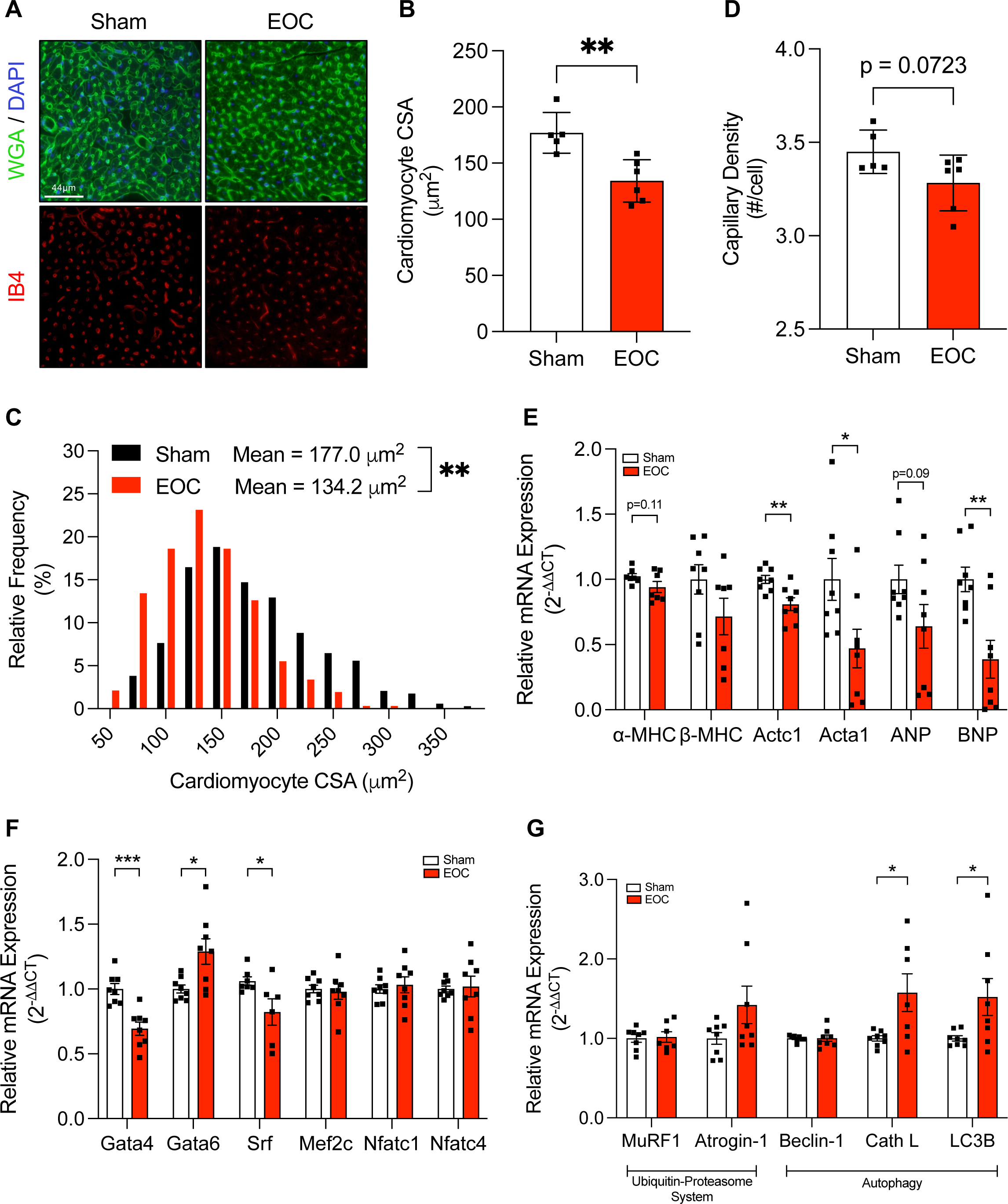
Ovarian tumor-bearing mice show cardiac atrophy, altered ventricle morphology, and upregulation of autophagy. (A) Representative LV cross sections from sham and EOC mice stained with wheat germ agglutinin (WGA; green) + DAPI (blue) (top) and isolectin B4 (red; bottom). Magnification 40x. (B) Cardiomyocyte cross-sectional area (CSA). (C) Capillary density. Data are means ± SD. (D) Histogram depiction of cardiomyocyte CSA distribution in sham and EOC mice analyzed on ranks using a Mann-Whitney test. (E) LV mRNA expression of _-myosin heavy chain (_-MHC; Myh6), β-myosin heavy chain (β-MHC; Myh7), _-cardiac actin (Actc1), _-skeletal actin (Acta1), atrial natriuretic peptide (ANP; Nppa), brain natriuretic peptide (BNP; Nppb). (F) LV mRNA expression of cardiac transcription factors; GATA binding protein 4 (Gata4), Gata6, Serum response factor (Srf), Myocyte enhancing factor 2C (Mef2c), Nuclear factor of activated T cells 1 (Nfatc1), Nfatc4. (G) LV mRNA expression of proteolytic pathway markers; MuRF1 (tripartite motif-containing 63), Atrogin-1 (F-box protein 32), Beclin-1, Cathepsin L (Cath L), microtubule-associated protein 1 light chain 3 beta (LC3B). mRNA data are means ± SEM. Significance determined by an unpaired student’s t-test.

We next measured the expression of genes critical for cardiac structure, myocyte contraction, and blood pressure regulation (Figure 5E). The expression of the predominant adult cardiac myosin heavy chain (MHC) isoform, lJ*-MHC*, tended to decrease (p=0.11) in EOC mice. Tumor-bearing mice also showed a downregulation of sarcomeric genes, C-cardiac actin (*Actc1*) and C-skeletal actin (*Acta1*). Expression of atrial (*ANP*) and brain natriuretic peptides (*BNP*), which function as cardioprotective hormones and blood pressure regulators, were also measured. *BNP* expression decreased in tumor-bearing mice and *ANP* expression showed a trending decrease as well (p=0.09). These findings reveal new evidence that ovarian cancer induces cardiomyocyte atrophy, altered ventricle morphology, and reduced expression of genes critical for normal cardiac structure and function.

### Cardiomyocyte atrophy due to ovarian cancer shows downregulation of transcription factors Gata4 and Srf

Cardiac transcription factors regulate cardiomyocyte structure and growth at the molecular level and are overexpressed in conditions of ventricular hypertrophy. Thus, we hypothesized that the expression of these transcription factors would decrease due to cancer. We observed divergent responses in mRNA expression of Gata transcription factors, where *Gata4* decreased and *Gata6* increased in EOC mice (Figure 5F). *Srf* also decreased in tumor-bearing mice. Transcription factors *Mef2c*, *Nfatc1*, and *Nfatc4* were expressed similarly between sham and EOC mice. Indeed, *Gata4* and *Srf* play an important role in the regulation of structural and cardioprotective genes (quantified in Figure 5E). These findings suggest that cardiac atrophy resulting from cancer could partly be due to attenuated *Gata4* and *Srf* expression.

### Autophagic, but not UPS, gene expression is elevated in ovarian tumor-bearing mice

Catabolic signaling is a pathogenic mediator of cardiac and skeletal muscle wasting in cancer. Thus, we measured the expression of several proteasomal markers of muscle wasting. Skeletal muscle atrophy is driven by upregulation of UPS-mediated protein degradation.^8^ However, *Murf1* and *Atrogin-1*, muscle-specific E3 ubiquitin ligases, did not increase in hearts of ovarian tumor-bearing mice (Figure 5G), suggesting that UPS-mediated protein degradation does not play a significant role in the development of cardiac atrophy with ovarian cancer.

Since hyperactivation of autophagic protein degradation promotes muscle wasting, we measured the expression of several genes involved in autophagy. Beclin-1, Cathepsin L, and LC3B are well-characterized markers of lysosomal activation and induction of autophagic degradation. We did not observe an increase in *Beclin-1* mRNA expression, an upstream regulator of autophagic sequestration. However, expression of *Cathepsin L*, a cysteine protease involved in lysosomal degradation, and *LC3B*, a marker of autophagosome formation, increased by ∼1.5-fold in tumor-bearing mice (Figure 5G), implicating autophagic activation as a likely contributing factor to cardiac atrophy due to cancer.

### Cardiac atrophy and dysfunction prevail in multiple tumor types

To characterize whether our findings in ovarian cancer are conserved across multiple tumor types, we assessed cardiac morphology and function in an orthotopic, syngeneic mouse model of advanced PDAC (Table S3). Pancreatic tumor-bearing mice similarly presented with a decrease in cardiac mass (∼8% compared to shams) and cardiomyocyte size (∼23% compared to shams; Figure 6A-D), with no change in the quantity of interstitial fibrosis (Figure S2B). Using invasive hemodynamics, PDAC mice showed an increase in heart rate compared to shams (Figure 6E). Maximum LV pressure (Table S3) and peak LV contraction and relaxation rates (Figure 6F) decreased in tumor-bearing mice, demonstrating new evidence that pancreatic cancer impairs cardiac function. Pancreatic tumor-bearing mice also showed decreases in mean arterial pressure (Figure 6G) and systolic and diastolic blood pressure (Figure 6H; Table S3). Together, these findings support that multiple tumor types cause cardiac atrophy, dysfunction, and systemic hypotension in advanced-stage cancer.

**Figure 6.**
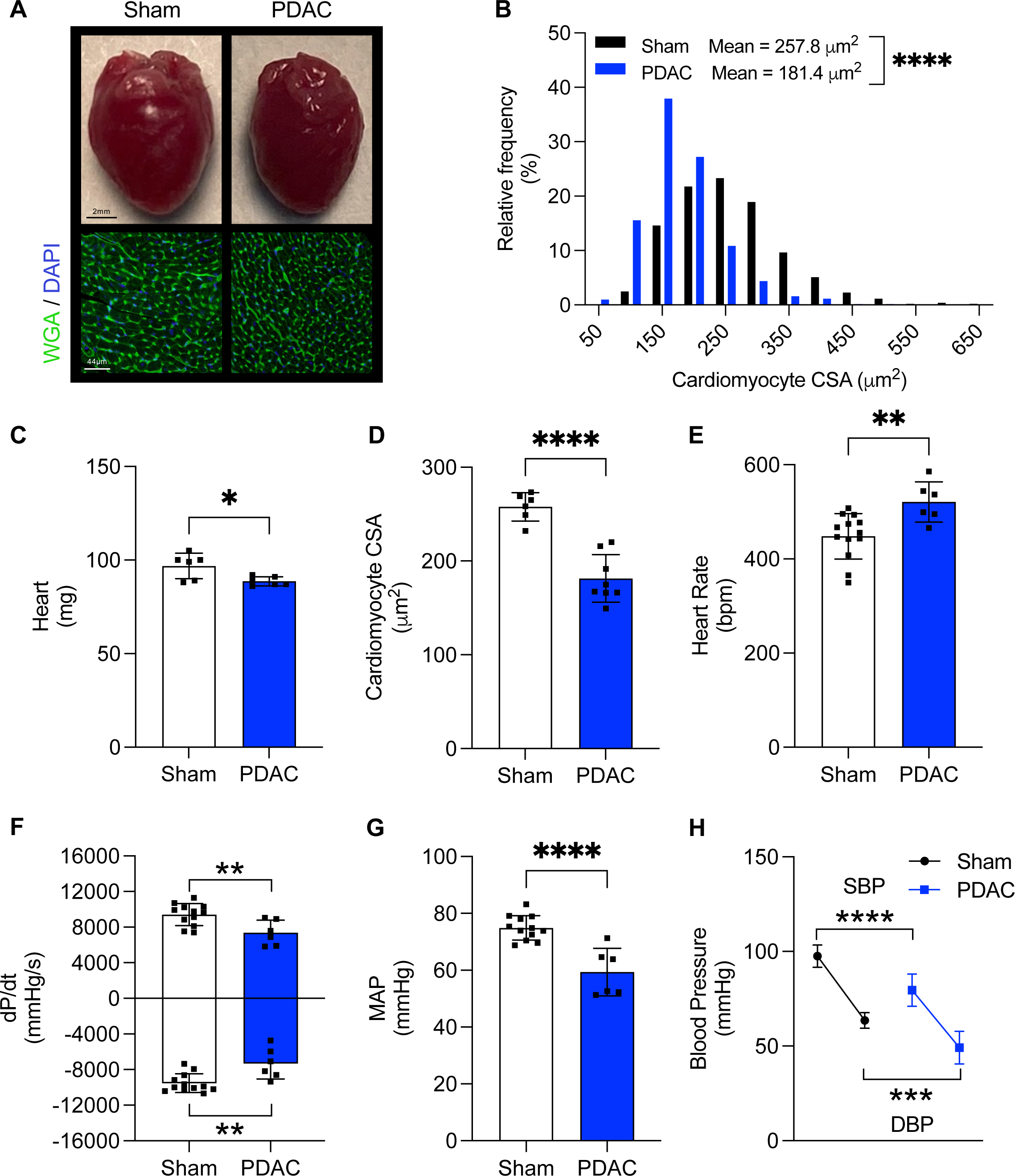
Cardiac atrophy, dysfunction, and hypotension in mice with advanced pancreatic cancer. (A) Representative gross morphology of hearts (top) and LV cross sections stained with wheat germ agglutinin (WGA; green) and DAPI (blue) (bottom) from sham and PDAC mice. Magnification 40x. (B) Histogram depiction of cardiomyocyte CSA distribution in sham and PDAC mice analyzed on ranks using a Mann-Whitney test. (C) Heart mass. (D) Cardiomyocyte CSA. (E) Heart rate. (F) dP/dt Max (top) and Min (bottom). (G) Mean arterial pressure (MAP). (H) Systolic (SBP; top) and diastolic blood pressure (DBP; bottom). Data are means ± SD. Significance determined by an unpaired student’s t-test.

### Human pancreatic cancer causes cardiac atrophy and induction of catabolic pathways

Since we established that murine tumors cause cardiac atrophy and dysfunction, we next sought to verify that this is consistent with human malignancy. Patient-derived pancreatic cancer xenografts were implanted orthotopically, and hearts were harvested for histological and molecular analyses. Cardiomyocyte size decreased (Figure 7A) and myocardial fibrosis tended to increase (p=0.08; Figure 7B, C) in tumor-bearing mice compared to shams. PDX mice showed attenuated α*-MHC* and increased *ß-MHC* gene expression (Figure 7D), indicating an MHC isoform shift that is consistent with a diseased cardiac phenotype. Tumor-bearing mice also showed a ∼2-fold decrease in *Actc1* expression (Figure 7D), supporting that human cancer alters the cardiac gene program.

**Figure 7.**
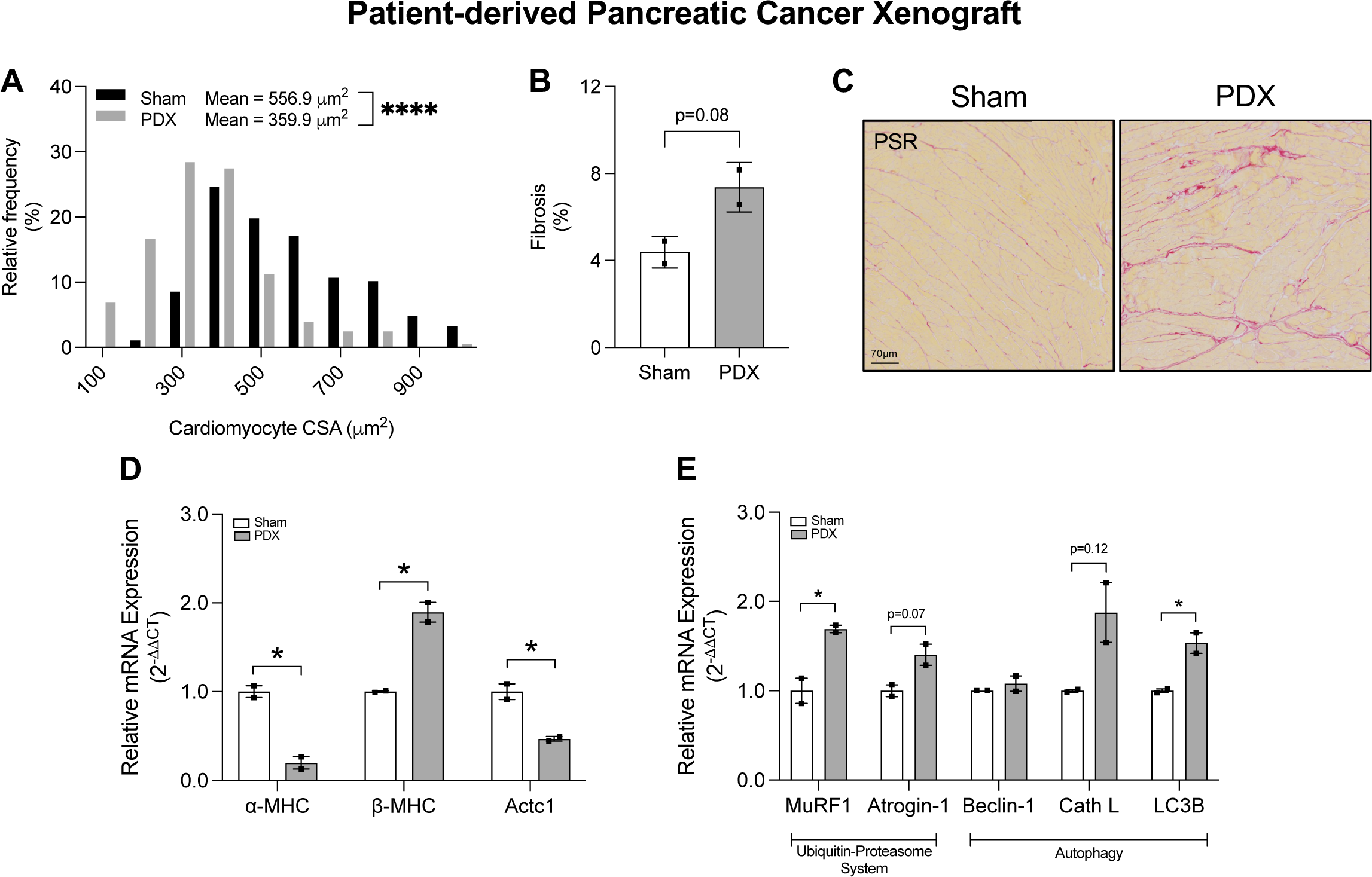
Pathological cardiac remodeling in a patient-derived xenograft model of pancreatic cancer. (A) Histogram depiction of cardiomyocyte CSA distribution in sham and PDX mice analyzed on ranks using a Mann-Whitney test. (B) Myocardial fibrosis. (C) Representative LV cross sections from sham and PDX mice stained with picrosirius red (PSR). Magnification 40x. (D) Cardiac mRNA expression of _-myosin heavy chain (_-MHC; Myh6), β-myosin heavy chain (β-MHC; Myh7), _-cardiac actin (Actc1). (E) Cardiac mRNA expression of proteolytic pathway markers; MuRF1 (tripartite motif-containing 63), Atrogin-1 (F-box protein 32), Beclin-1, Cathepsin L (Cath L), microtubule-associated protein 1 light chain 3 beta (LC3B). mRNA data are means ± SEM. Significance determined by an unpaired student’s t-test.

To elucidate the proteolytic mechanisms involved in cardiac atrophy, we measured markers of UPS and autophagy pathways (Figure 7E). In contrast to our observations in EOC mice showing no evidence of UPS pathway activation, PDX mice showed a 1.7-fold increase in *MuRF1* expression and a trend towards upregulation of *Atrogin-1* as well (p=0.07). Despite no differences in *Beclin-1* expression, *Cathepsin L* tended to increase (p=0.12) and *LC3B* increased by 1.5-fold in pancreatic tumor-bearing mice, consistent with autophagic induction. These findings demonstrate, for the first time, that both UPS and autophagy-mediated protein degradation are implicated in the development of cardiac atrophy in human pancreatic cancer.

## DISCUSSION

Here we investigated how advanced malignancy alters myocardial performance. While there has been significant focus on the effects of chemotherapy on cardiac health and cachexia, limited information exists on how the presence of malignancy alone impacts cardiac structure and function. We show that ovarian and pancreatic cancers induce cardiomyocyte atrophy (23-25% by cross-sectional area, 8-10% by mass), which occurs in parallel with decreases in LV contraction, relaxation, systemic hypotension, and severe exercise intolerance. Cardiac dysfunction was intrinsic to the heart and shown to be independent of neurohormonal input and hemodynamic loading status. The presence of a tumor promotes a substrate shift in cardiac metabolism to a fetal-like state that parallels metabolic dysfunction in failing hearts. In multiple tumor types, cancer alters the cardiac gene program to a molecular phenotype typical of patients with heart failure.^21^ Our data support the upregulation of autophagic and UPS signaling, which together contribute to the development of cardiac dysfunction and atrophy in advanced-stage cancers. Finally, we translated these impairments in cardiac structure using patient-derived xenografts of human cancer (Figure 8).

**Figure 8.**
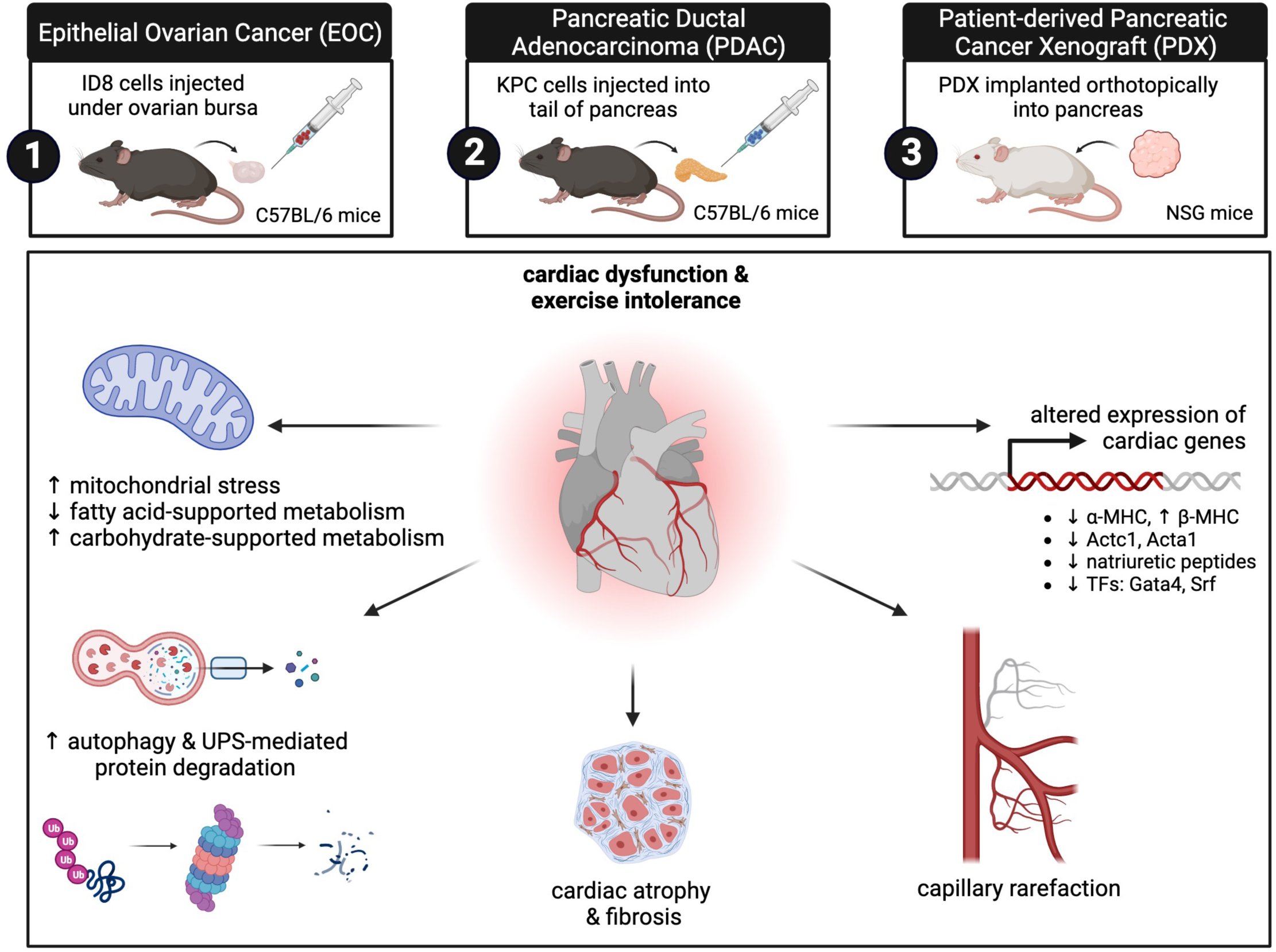
Cardiac atrophy, dysfunction, and metabolic impairments in advanced-stage cancers. TFs, transcription factors.

Cardiac dysfunction and atrophy have previously been observed in rodents^22,23^ and patients ^5^ with advanced cancer, yet, the molecular basis for these observations has received little investigation. Previous reports have demonstrated that cancer is associated with heightened activation of the sympathetic nervous system and renin-angiotensin-aldosterone system, increasing noradrenalin, renin, and aldosterone levels in rodents and patients with cancer.^22^ This is further supported by the ability of beta-adrenergic antagonists to reduce cardiac wasting and increase survival in rodents with cancer cachexia.^22^ Thus, neurohormonal activation is thought to be implicated in cancer-induced cardiomyopathy. We demonstrate new evidence that LV systolic and diastolic dysfunction persist in an *ex vivo* heart preparation – an assay that evaluates cardiac performance independent of extrinsic neurohormonal factors – supporting that cardiac dysfunction, at least in part, resides intrinsic to the myocardium in ovarian tumor-bearing mice.

Our findings show that the pathophysiology of cancer-related cardiac impairments is multifactorial. First, capillary density in the hearts of tumor-bearing mice tended to decrease. Since oxygen extraction in the myocardium is near maximum, capillary rarefaction causes ischemia, impaired cardiac metabolism, and severe dysfunction.^24^ Recently, the loss of vascular density was observed in skeletal muscle of tumor-bearing mice, which preceded the decline in muscle function and mass.^25^ Thus, endothelial cells that line the vessels of distant, non-metastatic organs are ideally situated as first responders to factors extruded from the primary tumor. We are the first to show that malignancy causes a loss of capillary density in the myocardium. Future temporal and mechanistic studies are warranted to determine whether myocardial endothelial cell loss precedes overt dysfunction and atrophic remodeling and to elucidate the signaling pathways that connect the tumor-heart axis.

Metabolic alterations are a central mechanism to both cardiovascular disease and cancer. Indeed, impairments in oxidative phosphorylation and mitochondrial biogenesis occur in anthracycline-induced cardiotoxicity.^26,27^ However, it is not known whether cancer itself directly perturbs cardiac metabolism. We show for the first time that cancer upregulates carbohydrate- and attenuating fatty acid-mediated mitochondrial respiration in the heart. The healthy heart relies predominantly on fatty acids as a fuel source, however, a greater reliance on glucose utilization occurs in fetal development and various cardiac disease states.^28^ While this metabolic shift is well documented in failing hearts, whether tumors can perturb substrate metabolism in the heart, to our knowledge, has not previously been shown. Recent evidence demonstrated that tumors could alter metabolic function of distant, non-metastatic organs, such as the liver. Wang et al. showed that tumor-secreted vesicles can suppress fatty acid oxidation and reprogram liver metabolism in the absence of hepatic metastases, which ultimately leads to liver dysfunction.^29^ Here we validate this concept, as our findings confirm that remote, advanced-stage tumors effectively reprogram cardiac metabolism to a fetal-like profile that is characteristic of decompensated heart failure. We propose that reduced myocardial perfusion in tumor-bearing mice leads to altered mitochondrial substrate selection from fatty acids to pyruvate oxidation. The degree to which this change in substrate selection contributes to cardiomyopathy during cancer is uncertain but may reflect an impairment in fatty acid oxidation that is matched by a partial compensation in pyruvate oxidation. Such increases in pyruvate oxidation in EOC contrast with decreases seen in hearts from a C26 colorectal cancer model, suggesting cardiac mitochondrial stress is either dependent on the cancer model or varies dynamically throughout cancer progression.^16^ Nonetheless, administration of the mitochondrial-targeting peptide SS-31, normalized contractile function in C26 tumor-bearing mice, demonstrating that mitochondrial stress is critically involved in the detrimental effects that cancer has on the cardiovascular system.

Proteolytic pathways play important regulatory roles in protein turnover in the healthy heart,^30^ yet the role of each system in maintaining cardiac structure and function remains controversial. Metabolic remodeling in energy-depleted states, such as cachexia, induces activation of the UPS and autophagic pathways to promote protein catabolism in muscle tissue. Indeed, hyperactivation of the UPS has been implicated in skeletal muscle atrophy, however its role in cancer-induced cardiac atrophy has been debated. Zhou et al. showed that colon-26 cancer cachexia caused cardiac atrophy via activation of the UPS, which was abolished by pharmacological type IIB activin receptor pathway blockade.^31^ Conversely, others have shown that autophagic signaling plays a larger role in cancer-induced cardiac atrophy, where products of autophagic degradation are recycled to support increased metabolic demand.^16,32^ Cosper et al. identified that cardiac atrophy is associated with an increase in autophagy, but not UPS activity in colon-26 cancer cachexia, which they attributed to the fact that basal UPS activity is already high in the normal heart.^32^ Our findings support a role for both UPS- and autophagy-mediated protein degradation in hearts of mice with advanced-stage cancer, with slight variation that is possibly the result of speciation or malignancy type/stage. While mice bearing ovarian tumors only showed an increase in markers of autophagy, mice bearing humanized pancreatic tumors showed evidence that both pathways are implicated in the etiology of cardiac atrophy. The contributions of each pathway in myocardial protein catabolism may depend on the tumor type, stage, and varying composition of factors derived from the tumor, thus warranting further investigation.

In the current study, cardiac atrophy, functional impairments, and arterial hypotension in tumor-bearing mice were associated with altered expression of cardiac transcription factors, *Gata4* and *Srf*. These factors support basal transcription of cardiac genes and play important roles in ventricular remodeling and cardioprotection in response to various stimuli.^12^ GATA4 is one of the earliest expressed transcription factors in the developing myocardium and its targets include α- and ß-MHC, natriuretic peptides, and other sarcomeric genes.^33^ Thus, pathological alterations in the cardiac gene program in tumor-bearing mice – decreased α*-MHC* and natriuretic peptides, increased *ß-MHC* expression – could be explained, at least in part, by a decrease in *Gata4* expression. Others have shown that in *Gata4*-deficient conditions, the upregulation of *Gata6* is capable of compensating for the lack of *Gata4* to maintain cardiac gene expression.^34^ However, our data show that cancer decreases *Gata4* expression, which is associated with attenuations of several cardiac genes, that cannot be mitigated by an upregulation in endogenous *Gata6* expression. Ovarian tumor-bearing mice also showed decreased expression of *Srf* – a transcriptional activator of cardiac sarcomeric genes. The loss of *Srf* in the adult heart impairs LV contractility, relaxation, and decreases wall thickness, which is associated with reduced *Actc1* and *Acta1* expression.^35^ Our data support a critical role for *Srf* in maintaining expression of cardiac and skeletal muscle actin, myocyte structure, and function, which is attenuated in advanced cancer. Previous literature shows the importance of transcription factors in maintaining cardiac health,^13^ but we do not yet understand how decreases in these cardiac transcription factors occur in the presence of a tumor.

In the past decade, there has been growing interest in understanding how cancer therapies contribute to short-term cardiac complications and influence long-term quality of life and morbidity in cancer patients. Many anti-cancer therapies have negative consequences on the cardiovascular system, which can limit their administration or dosing, severely impact patient quality of life and recovery, or lead to chronic cardiovascular conditions, where patients require additional medications to manage irreversible cardiac damage.^3^ Cardiac atrophy and dysfunction have previously been demonstrated in therapy-naïve cancer.^22,23,36^ However, preclinical work in this area has predominantly used ectopic cancer models to investigate how cachexia impacts the heart. While this work has been fundamental for characterizing the systemic effects of cachexia on the cardiovascular system, the use of orthotopic cancer models, such as those employed in the current study, provide the ability to recapitulate the tumor microenvironment that occurs in human cancers and conduct investigations that improve clinical translation. Further, the use of patient-derived xenograft models provide the opportunity to conduct mechanistic investigations that elucidate the molecular pathways driving cardiac dysfunction and therapeutic studies that explore cardioprotective strategies in human cancer. Using clinically relevant cancer models, we highlight that cardiac atrophy is associated with metabolic and molecular profiles that are consistent with heart failure.^21^ These findings demonstrate translational evidence that pathological cardiac remodeling is a feature of human cancer and should be considered in the management of this patient population. This is particularly important for patients undergoing cardiotoxic cancer therapy and patients with CVD risk factors or pre-existing heart disease, where direct consequences of cancer on the heart may be a compounding detriment to cardiovascular health.

## STUDY LIMITATIONS

There are several limitations to this study. Male and female mice were used to evaluate how KPC-based pancreatic cancer impacts cardiac function. Sex differences have previously been shown in the severity and underlying mechanisms of cardiac pathology in cancer cachexia.^32^ However, we did not stratify data by sex to discern potential differences, as the primary objective of this research was to investigate cardiac dysfunction and atrophy across various tumor types. Next, we measured mRNA expression of catabolic pathway markers, however, direct evidence of autophagic and UPS activity or myofibrillar protein degradation were not measured. Others have shown that markers of autophagy and the UPS are transcriptionally upregulated in conditions of muscle catabolism, supporting that this indicates proteolytic pathway activation,^37,38^ but further studies are needed to assess autophagic flux, organelle, and proteasome function in depth.

## CONCLUSIONS

Here we demonstrate new insight into the cardiac structural and functional impairments from multiple types of advanced-stage cancers. Together, our data highlight that the presence of a tumor elicits an energetic stress in the myocardium by decreasing vascularization and perturbing cardiac metabolism. Consequently, this promotes autophagy and UPS activation, and alterations in the cardiac gene program that parallel a diseased molecular phenotype, ultimately leading to cardiac atrophy and failure. These observations were validated in a patient-derived xenograft model, providing translational evidence of cardiac pathology induced by human cancer. Our findings confirm that cancer is a direct insult on cardiac physiology, highlighting the importance of managing cardiac health in cancer patients.

## METHODS

### Animals

C57BL/6 (Charles River Laboratories) and NOD-SCID IL2 receptor gamma chain knockout (NSG) mice (Jackson Laboratories) were housed in a temperature-controlled, 12h light:dark cycle and given food and water *ad libitum*. NSG mice were housed in virus-free conditions. All procedures were approved by the Animal Care Committee at the University of Guelph and in compliance with Canadian Council on Animal Care guidelines or guidelines from the Virginia Commonwealth University Institutional Animal Care and Use Committee.

### Reagents and Cell Lines

Spontaneously transformed murine ovarian surface epithelial cells (ID8; generously donated by Drs. K. Roby and P. Terranova, Kansas State University, Manhattan, KS) were cultured in Dulbecco’s Modified Eagle Medium (Wisent Inc.) with 10% fetal bovine serum and 1% antibiotic/antimycotic (Wisent, Inc.). KPC cells (generously donated by Dr. Steven Gallinger, University of Toronto, Toronto, ON) were cultured in RPMI (Wisent Inc.) with 10% fetal bovine serum, 1% antibiotic/antimycotic, 1% sodium pyruvate (Wisent Inc).

### Epithelial Ovarian Cancer Mouse Model

In female C57BL/6 mice at 9-10 weeks of age, ovarian tumors were induced as an orthotopic, syngeneic mouse model of EOC. Briefly, a dorsal incision was made on isoflurane-anesthetized mice and 1.0×10^6^ ID8 cells in 5uL sterile saline were injected under the bursa of the left ovary. Sham mice followed the same procedure with injection of only sterile saline. In this model, large primary tumors form ∼8.5 weeks after tumor cell injection, followed by the development of abdominal ascites and secondary peritoneal lesions, at which point the model has disease characteristics that correspond to a clinical profile of stage III EOC.^39^ To evaluate cardiac health in advanced-stage ovarian cancer, tumors were allowed to develop for approximately 13 weeks after cell implantation, then *in vivo* experiments were performed and tissues were collected.

### Pancreatic Ductal Adenocarcinoma Mouse Model

At 9-10 weeks of age, pancreatic tumors were induced as an orthotopic, syngeneic, immunocompetent mouse model of PDAC. Briefly, male and female C57BL/6 mice were anesthetized with isoflurane, a dorsal incision was made and 1.0×10^5^ KPC cells were injected into the tail of the pancreas. Sham mice followed the same procedure with injection of only sterile saline. To evaluate cardiac physiology in advanced-stage pancreatic cancer, tumors were allowed to develop for approximately 4 weeks after cell implantation, then cardiac hemodynamic and histological analyses were performed.

### Patient-derived Xenograft Model

A viable 2×2mm portion of tissue was isolated from a surgically resected primary pancreatic cancer specimen with minimal ischemia time. Tissue was implanted subcutaneously into male (NSG) mice (Jackson Laboratory). Xenografts were allowed to grow to a maximum diameter of 1.5cm before passage. Herein, we defined passage as explantation of a pancreatic cancer xenograft and orthotopic implantation into the pancreas of a new host. Xenografts were allowed to grow for approximately 20 weeks, then mice were euthanized and hearts were collected for histological and molecular analyses.

### Voluntary Wheel Running

Mice were individually housed in cages with access to an exercise wheel and a cycle computer (VDO) was used to record running distance. Mice were given a 24-hour acclimation period, then data recorded for two consecutive 24-hour periods were averaged.

### Echocardiography

Mice were anesthetized with an isoflurane/oxygen mix (2%:100%) and body temperature maintained at 37.2°C-37.5°C. M-Mode echocardiography was performed on the left ventricle (LV) using the Vevo2100 system (VisualSonics Inc.) with a MS550D transducer.

### Invasive Hemodynamics

Mice were anesthetized with an isoflurane/oxygen mix (2.5%:100%) and body temperature maintained at 37.2°C-37.5°C. The right carotid artery was isolated and a 1.2Fr pressure catheter (Transonic Scisense Inc.) was inserted and advanced into the LV. Hemodynamic signals were sampled at a rate of 2kHz and analysis performed with Spike2 software (Cambridge Electronic Design, Ltd.).

### Langendorff

Mice were anesthetized with an isoflurane/oxygen mix (2.5%/100%). Hearts were excised and rinsed in ice-cold phosphate buffered saline (PBS). The aorta was mounted on a 20-gauge cannula and hearts were retrograde-perfused at a constant pressure of 70-75mmHg with warmed (37°C) and oxygenated (95% O_2_:5% CO_2_) Krebs-Henseleit buffer (pH 7.4), containing in (mmol/L) 118 NaCl, 4.7 KCl, 1.2 MgSO_4_, 1.2 KH_2_PO_4_, 0.5 C_3_H_3_NaO_3_, 0.05 EDTA, 11 glucose, and 2 CaCl_2_. After stable perfusion rates were achieved, the left atrial appendage was removed, and a deflated balloon connected to a pressure transducer was inserted into the LV to record pressure. The balloon was inflated in a stepwise manner (∼2 mmHg change in end diastolic pressure; EDP) until a set EDP of 5-8 mmHg was reached. The maximal rates of LV contraction (dP/dt Max) and relaxation (dP/dt Min) and maximal LV pressure were obtained from raw pressure tracings using Spike2 software (Cambridge Electronic Design Ltd.).

### Preparation of permeabilized muscle fibres for mitochondrial respiration and H_2_O_2_ (mH_2_O_2_) emission

This technique was adapted from previous methods described elsewhere.^40^ Briefly, the heart was removed and placed in ice-cold BIOPS, containing (in mM): 50 MES hydrate, 7.23 K_2_EGTA, 20 imidazole, 0.5 dithiothreitol, 20 taurine, 5.77 ATP, 15 PCr and 6.56 MgCl2·6H2O (pH 7.2). The LV was isolated and gently separated along the longitudinal axis to form fibre bundles (PmFBs), which were blotted and weighed (0.8-2.1mg wet weight) in 1.5 mL of tared, prechilled BIOPS to ensure PmFBs remained relaxed and hydrated. These bundles were treated with 40μg/mL saponin in BIOPS on a rotor for 30 minutes at 4°C to selectively permeabilize the cell membrane. PmFBs that were used for mH_2_O_2_ were also treated with 35μM 2,4-dinitrochlorobenzene (CDNB) during the permeabilization step to deplete glutathione and allow for detectable rates of mH_2_O_2_.^41^ All PmFBs were then washed in Buffer Z (containing in mM: 105 K-MES, 50 KCl, 10 KH_2_PO_4_, 5 MgCl_2_·6 H_2_O, 1 EGTA and 5 mg/ml BSA; pH 7.2) on a rotator for 15 minutes at 4°C to remove the cytoplasm.

High-resolution O_2_ consumption measurements were conducted in 2mL of respiration media (Buffer Z) using the Oxygraph-2k (Oroboros Instruments, Austria) while stirring at 750rpm at 37°C. Buffer Z contained 20mM creatine to saturate mtCK (mitochondrial creatine kinase) and promote phosphate shuttling through mtCK.^42^ For carbohydrate-supported ADP-stimulated respiratory kinetics, 5mM pyruvate and 2mM malate were added as Complex I substrates (via generation of NADH to saturate electron entry into Complex I) followed by titrations of submaximal ADP (25µM, 100µM, 500µM and 1000µM) and maximal ADP (5000µM). 10mM glutamate was also added to the assay media to further saturate Complex I with NADH. Last, cytochrome *c* was added to test for outer mitochondrial membrane integrity and succinate (20mM) was added to saturate electron entry into Complex II with FADH_2_. For fatty acid-supported ADP-stimulated respiratory kinetics, 5mM L-carnitine, 0.02mM Palmitoyl-CoA and 0.5mM malate were added as Complex I, II, and electron transport chain (ETC) substrates (via generation of NADH and FADH_2_ from ß-oxidation) followed by titrations of submaximal ADP (100µM and 300µM) and maximal ADP (500µM). Cytochrome *c* was added to test for outer mitochondrial membrane integrity and succinate (20mM) was added to saturate electron entry into Complex II. All experiments were completed in the presence of 5µM BLEB in the assay medium to prevent ADP-induced contraction of PmFBs.^42^

mH_2_O_2_ was determined spectrofluorometrically (QuantaMaster 40, HORIBA Scientific) in a quartz cuvette with continuous stirring at 37°C, in 1mL of Buffer Z supplemented with 10μM Amplex Ultra Red, 0.5U/mL horseradish peroxidase, 1mM EGTA, 40U/mL Cu/Zn-SOD1, 5μM BLEB, and 20mM Cr. Buffer Z contained (in mM) 105 K-MES, 30 KCl, 10 KH_2_PO_4_, 5 MgCl_2_·6H_2_O, and 1 EGTA and 5 mg/mL BSA (pH 7.2). State II mH_2_O_2_ (maximal emission in the absence of ADP) was induced using the Complex I-supporting substrates (NADH) pyruvate (10mM) and malate (2mM) as described previously.^43^ Following the induction of state II mH_2_O_2_, a titration of ADP was employed to progressively attenuate mH_2_O_2_ as occurs when membrane potential declines during oxidative phosphorylation. The rate of mH_2_O_2_ emission was calculated from the slope (F/min) using a standard curve established with the same reaction conditions and normalized to fibre bundle wet weight.

### Western Blotting

LV tissue was homogenized in ice-cold buffer containing (in mM) 20 Tris, 150 NaCl, 1 EDTA, 1 EGTA, 2.5 Na_4_O_7_P_2_, 1% Triton X-100 with PhosSTOP inhibitor tablet (MilliporeSigma) and protease inhibitor cocktail (MilliporeSigma). 7μg of protein, measured by a BCA protein assay kit (ThermoFisher Scientific Inc.) was subjected to 12% SDS-PAGE followed by transfer to polyvinylidene difluoride membrane. Membranes were blocked with Odyssey blocking buffer (Li-COR) and immunoblotted overnight (4°C) with rodent OXPHOS cocktail monoclonal antibody (Abcam; ab110413, 1:1000) to detect ETC proteins. Membranes were washed and incubated with an infrared fluorescent secondary antibody (LI-COR, 1:20,000). Immunoreactive proteins were detected by infrared imaging and quantified by densitometry (ImageJ, NIH). Images were normalized to Amido Black total protein stain (MilliporeSigma).

### Histological Analysis

LV tissue was fixed in 10% neutral buffered formalin and paraffin-embedded cross sections (5 µm) were stained with either Picrosirius Red (Sigma Aldrich) to visualize interstitial fibrosis or wheat germ agglutinin AlexaFluor^TM^ 488 Conjugate (ThermoFisher Scientific Inc.) and isolectin GS-IB4 AlexaFluor^TM^ 568 Conjugate (ThermoFisher Scientific Inc.) to assess myocyte cross-sectional area (CSA) and capillary density, respectively. DAPI (Invitrogen) was used as a fluorescent nuclear counterstain. Images were acquired using an Olympus FSX 100 light microscope analyzed with CellSens software (Olympus) to quantify interstitial fibrosis, or ImageJ software (NIH) to quantify CSA and capillary density.

### Quantitative Reverse Transcription (RT)-PCR

Total RNA was isolated using a TRIzol (Invitrogen) and RNeasy (Qiagen) hybrid protocol. Briefly, LV tissue was homogenized in TRIzol reagent according to the manufacturer instructions. The RNA mixture was transferred to a RNeasy spin column (Qiagen) and processed according to the RNeasy Kit instructions. RNA was quantified spectrophotometrically at 260nm using a NanoDrop (ND1000, ThermoFisher Scientific Inc.). RNA was reverse transcribed for qPCR using a High-Capacity cDNA Reverse Transcription Kit (Applied Biosystems). Quantitative PCR reactions were performed on a CFX Connect Real-Time PCR System (Bio-Rad Laboratories, Ltd.) using SsoAdvanced Universal SYBR Green Supermix (Bio-Rad Laboratories, Ltd.). Primer sequences are provided in Table S1. Data were normalized using *Rpl32* (ribosomal protein L32) as a reference gene. Statistical analysis of mRNA expression was performed on linear data using the 2^-ΔΔCT^ method.

### Statistical Analysis

Graphical and statistical analyses were completed using Prism 10 (GraphPad). Data are presented as mean ± SD unless otherwise stated. Gaussian distribution was tested using a Shapiro-Wilk test for normality. Unpaired t-tests were used to determine group differences, where appropriate. The distribution of CSA was analyzed on ranks using a Mann-Whitney test. Mitochondrial respiration and H_2_O_2_ emission were analyzed by two-way ANOVA with Šídák’s multiple comparisons test. Differences were considered significant at p<0.05. *p<0.05, **p<0.01, ***p<0.001, ****p<0.0001.

## ACKNOWLEDGEMENTS

Figure 8 created with Biorender.com.

