## Supplementary material for "Cardiac atrophy, dysfunction, and metabolic impairments: a cancer-induced heart failure phenotype": Data Supplement

**SUPPLEMENTAL DATA**

**Table S1. List of primer sequences used for reverse transcription qPCR.**

| ***Gene*** | ***Forward Primer (5’-3’)*** | ***Reverse Primer (5’-3’)*** |
| --- | --- | --- |
| *Myh6* | CACCAACAACCCATACGACTAC | TCAGCACATCAAAGGCACTATC |
| *Myh7* | AGATGGCTGGTTTGGATGAG | TTGGCCTTGGTCAGAGTATTG |
| *Nppa* | GGGTAGGATTGACAGGATTGG | TTCCTCCTTGGCTGTTATCTTC |
| *Nppb* | GGGAGAACACGGCATCATT | CCCAGCGGTGACAGATAAAG |
| *Acta1* | CCCAAAGCTAACCGGGAGAAG | GACAGCACCGCCTGGATAG |
| *Actc1* | CCCCGTCCATCAGAGAGCTA | TGGGTTCTGTAGGCGTGCTA |
| *Mef2c* | GTGGTTTCCGTAGCAACTCCTAC | GGCAGTGTTGAAGCCAGACAGA |
| *Gata4* | GCCTCTATCACAAGATGAACGGC | TACAGGCTCACCCTCGGCATTA |
| *Gata6* | ATGCGGTCTCTACAGCAAGATGA | CGCCATAAGGTAGTGGTTGTGG |
| *Srf* | CTCACCTACCAGGTGTCGGAAT | CTGCTGACTTGCATGGTGGTAG |
| *Nfatc1* | GGTGCCTTTTGCGAGCAGTATC | CGTATGGACCAGAATGTGACGG |
| *Nfatc4* | GATCGAGGTACAGCCTAGAGCA | GCAGAGTCAATGGCTTCTCACTG |
| *MuRF1* | TACCAAGCCTGTGGTCATCCTG | ACGGAAACGACCTCCAGACATG |
| *Atrogin-1* | CTTCTCGACTGCCATCCTGGAT | TCTTTTGGGCGATGCCACTCAG |
| *Becn1* | CAGCCTCTGAAACTGGACACGA | CTCTCCTGAGTTAGCCTCTTCC |
| *Cath L* | GGAAAATGGAGGTCTGGACTCG | GTGTCATTAGCCACAGCGAACTC |
| *LC3B* | GTCCTGGACAAGACCAAGTTCC | CCATTCACCAGGAGGAAGAAGG |
| *Rpl32* | GCCTCTGGTGAAGCCCAAG | TTGTTGCTCCCATAACCGATGT |

*Myh6*, α-myosin heavy chain; *Myh7*, ß-myosin heavy chain; *Nppa*, natriuretic peptide A (also known as atrial natriuretic peptide, ANP); *Nppb*, natriuretic peptide B (also known as brain natriuretic peptide, BNP); *Acta1*, α-skeletal actin; *Actc1*, α-cardiac actin; *Mef2c*, myocyte enhancing factor 2C; *Gata4*, GATA binding protein 4; *Gata6*, GATA binding protein 6; *Srf*, Serum response factor; *Nfatc1*, nuclear factor or activated T cells 1; *Nfatc4*, nuclear factor or activated T cells 4; *Trim63*, tripartite motif-containing 63 (also known as MuRF1); *Fbxo32*, F-box protein 32 (also known as MAFbx, Atrogin-1); *Becn1*, Beclin-1; *Ctsl*, Cathepsin L (also known as Cath L); *Map1lc3b*, microtubule-associated protein 1 light chain 3 beta (also known as LC3B); *Rpl32*, ribosomal protein L32.

**Table S2. Comparison of morphometric parameters between sham and ovarian cancer mice.**

|  | **Sham** | **EOC** | **p-value (summary)** |
| --- | --- | --- | --- |
| Tumour-free body weight (g) | 20.9 ± 1.4 | 19.8 ± 2.3 | 0.014 (*) |
| Ovary / tumour (mg) | 7.3 ± 1.6 | 251.0 ± 86.1 | <0.0001(****) |
| Spleen (mg) | 69.1 ± 10.0 | 206.2 ± 88.4 | <0.0001(****) |
| Kidney (mg) | 123.1 ± 12.5 | 114.6 ± 19.5 | 0.105 (ns) |
| Liver (mg) | 964.8 ± 174.2 | 997.8 ± 174.2 | 0.807 (ns) |
| TA (mg) | 39.6 ± 3.9 | 28.7 ± 6.8 | <0.0001 (****) |
| EDL (mg) | 7.2 ± 1.3 | 5.9 ± 1.7 | 0.014 (*) |
| Soleus (mg) | 7.3 ± 1.0 | 5.9 ± 1.5 | 0.001 (**) |
| Hematocrit (%) | 43.1 ± 3.1 | 27.5 ± 6.7 | <0.0001 (****) |

Values are mean ± SD. n=12-36 per group. EOC, epithelial ovarian cancer; TA, tibialis anterior; EDL, extensor digitorum longus. Significance determined by an unpaired student’s t-test; ns, not significant.

**Table S3. Comparison of morphometric and hemodynamic parameters between sham and pancreatic cancer mice.**

|  | **Sham** | **PDAC** | **p-value (summary)** |
| --- | --- | --- | --- |
| *Morphometric Parameters*  (n=6-8) | | | |
| Tumour-free body weight (g) | 23.8 ± 3.0 | 22.5 ± 1.7 | 0.535 (ns) |
| Pancreas / tumour (mg) | 167.3 ± 23 | 860.9 ± 385 | 0.0011 (**) |
| Heart (mg) | 96.8 ± 6.8 | 88.7 ± 2.4 | 0.0197 (*) |
| Cardiomyocyte CSA (μm) | 257.8 ± 86.3 | 179.6 ± 62.3 | p<0.0001 (****) |
| *Invasive Hemodynamic Parameters*  (n=6-12 per group) | | | |
| LVP Max (mmHg) | 102.9 ± 10.4 | 90.2 ± 15.2 | 0.0245 (*) |
| End diastolic pressure (mmHg) | 4.3 ± 2.2 | 5.1 ± 3.5 | 0.5786 (ns) |
| dP/dt Max (mmHg/s) | 9410 ± 1232 | 7368 ± 1416 | 0.0061 (**) |
| dP/dt @LVP40 (mmHg/s) | 8829 ± 1074 | 7183 ± 1318 | 0.0116 (*) |
| dP/dt Min (mmHg/s) | -9523 ± 1047 | -7320 ± 1740 | 0.0038 (**) |
| Tau (Glantz) (ms) | 8.9 ± 1.3 | 9.9 ± 3.4 | 0.8788 (ns) |
| Heart Rate (bpm) | 448 ± 48 | 521 ± 43 | 0.0056 (**) |
| Systolic BP (mmHg) | 97.5 ± 5.9 | 79.5 ± 8.6 | p<0.0001 (****) |
| Diastolic BP (mmHg) | 63.6 ± 4.1 | 49.2 ± 8.6 | 0.0002 (***) |
| MAP (mmHg) | 74.9 ± 4.3 | 59.3 ± 8.4 | p<0.0001 (****) |

Values are mean ± SD. PDAC, pancreatic ductal adenocarcinoma; LVP Max, maximum left ventricle pressure; dP/dt Max, maximum rate of change of pressure during systole; dP/dt @LVP40, rate of change of pressure at 40mmHg during systole; dP/dt Min, maximum negative rate of change of pressure during diastole; Tau, relaxation time constant; BP, blood pressure; MAP, mean arterial pressure. Significance determined by an unpaired, student’s t-test; ns, not significant.

**FIGURE LEGENDS**

**Figure S1. Mitochondrial oxygen consumption.** (A) Representative western blot images for ETC protein content (left) normalized to Amido Black stain (right). Carbohydrate ADP-stimulated Respiration; (B) Pyruvate & Malate, (C) Δ Glutamate, and (D) Δ Succinate. Fatty acid-ADP-stimulated Respiration; (E) L-Carnitine + Palmitoyl CoA and (F) Δ Succinate. Data are means ± SD. Significance determined by an unpaired student’s t-test.

**Figure S2. Quantification of myocardial fibrosis.** Left ventricular fibrosis quantity in (A) EOC and (B) PDAC mice compared to shams. Data are means ± SD. Determined by an unpaired student’s t-test.
