## Supplementary figures and images for "Cardiac atrophy, dysfunction, and metabolic impairments: a cancer-induced heart failure phenotype"

### Figure S1

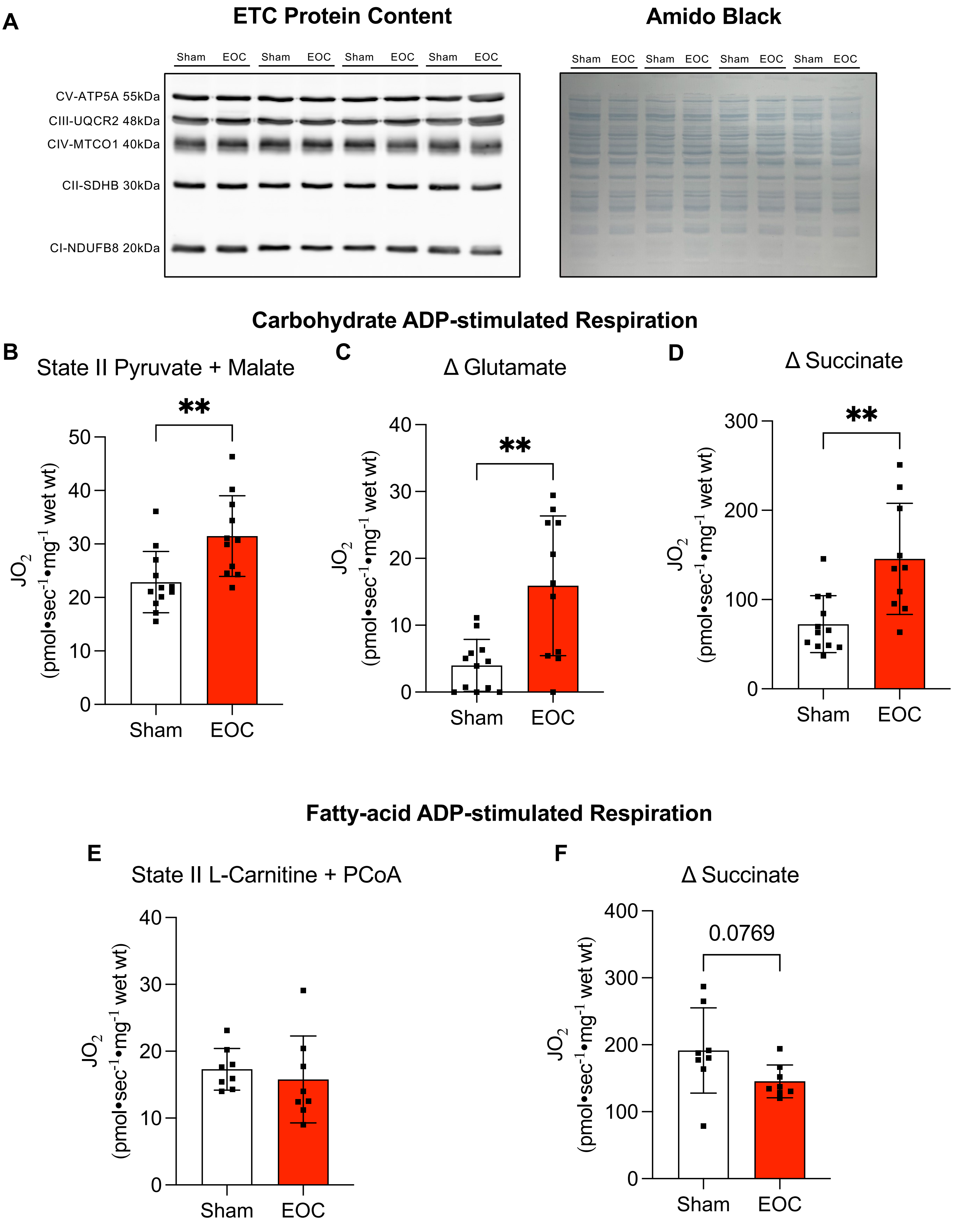

### Figure S2

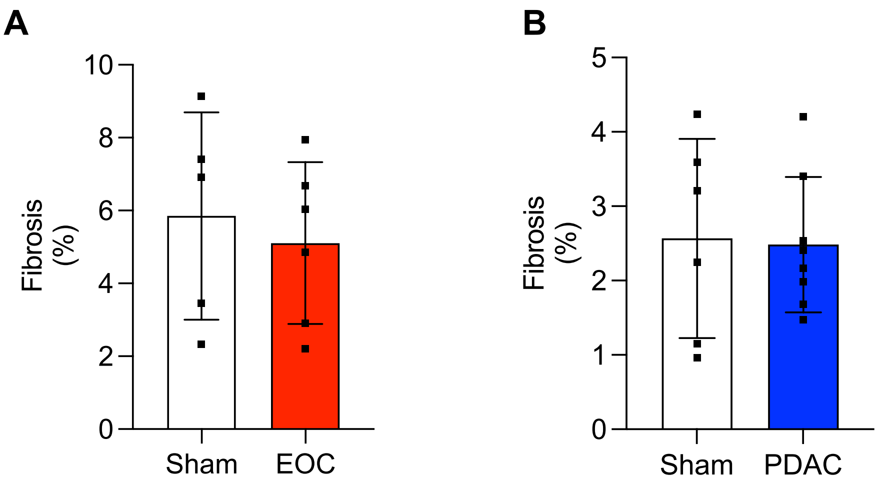


**Figure S2.**
